# Combination Ad26.RSV.preF/RSV preF protein vaccine induces superior humoral, cellular, and protective immunity compared with individual vaccine components in preclinical models

**DOI:** 10.1101/2022.09.23.509196

**Authors:** Eirikur Saeland, Leslie van der Fits, Renske Bolder, Marjolein Heemskerk-van der Meer, Joke Drijver, Yolinda van Polanen, Cornelis Vaneman, Lisanne Tettero, Freek Cox, Jan Serroyen, Matthew J. Jorgenssen, Johannes P.M. Langedijk, Hanneke Schuitemaker, Benoit Callendret, Roland C. Zahn

## Abstract

Respiratory syncytial virus (RSV) is a leading cause of severe respiratory disease for which no licensed vaccine is available. We have previously shown that a prefusion (preF) conformation– stabilized RSV F protein antigen and an adenoviral vector encoding RSV preF protein (Ad26.RSV.preF) are immunogenic and protective in animals when administered as single components. Here, we evaluated a combination of the 2 components. Strong induction of both humoral and cellular responses was shown in RSV-naïve and pre-exposed mice and pre-exposed African green monkeys (AGMs). Both components of the combination vaccine contributed to humoral immune responses, while the Ad26.RSV.preF component was the main contributor to cellular immune responses in both mice and AGMs. Immunization with the combination elicited superior protection against RSV A2 challenge in cotton rats. These results demonstrate the advantage of a combination vaccine and support further clinical development.

## Introduction

Respiratory syncytial virus (RSV) infections are a common cause of severe respiratory disease in older and immunocompromised adults,^1^ children,^2^ and those with certain chronic cardiac or pulmonary comorbidities.^3^ Annually, RSV is estimated to cause >60 million acute respiratory infections (ARIs) in adults and children worldwide.^4^ A recent prospective global study found that RSV causes disease burden in infected adults similar to or worse than influenza, as measured by clinical symptom scores and medical resource utilization.^5^ Older adults are at particularly high risk for severe RSV-mediated disease; in the United States alone, there are an estimated 177,000 hospitalizations and 14,000 deaths due to RSV in adults aged ≥65 years.^6^ The high RSV-associated morbidity in older adults may result from waning or impaired immune responses with age, referred to as “immune senescence”.^7, 8^ Given the high morbidity and significant disease burden of RSV in older adults, an effective prophylactic vaccine is needed to protect this population.

The RSV fusion glycoprotein (RSV F) is an attractive vaccine antigen because it is the main target of RSV virus–neutralizing antibodies (VNAs) in human sera.^9, 10^ RSV F fuses the viral and host cell membranes by irreversible protein refolding from the labile prefusion (preF) conformation to the stable postfusion (postF) conformation.^11^ RSV postF–based vaccines have shown limited or no protective efficacy in clinical trials,^12^ possibly because human RSV virus– neutralizing antibodies (VNAs) are primarily formed against RSV preF protein.^10, 13^ RSV preF– based vaccine development has long been hindered by the instability of RSV preF; however, recent efforts to stabilize the protein in the preF conformation using mutations identified by structure-based design have been successful.^11, 14^ Compared with postF-based vaccines, RSV preF–based vaccines have demonstrated superior efficacy and immunogenicity in preclinical studies.^11,14–16^

Although there is no commonly agreed-upon correlate of protection (CoP) against RSV infection, there is a wealth of literature establishing the importance of VNAs for preventing RSV infection.^17^ VNAs and their correlation with protection are typically measured in the serum; however, some evidence exists that mucosal RSV-specific immunoglobulins (Igs), particularly IgA, are significantly associated with reduced RSV infection rates and disease severity.^18, 19^ Prior studies also suggest that CD4+ and CD8+ T cells play an important role in the immune response to RSV infection.^17, 20^ Airway CD8+ T cells have been shown to play a protective role against RSV disease in humans^21^ and mice,^22, 23^ whereas CD4+ follicular helper T cells support antibody production, class switching, and memory B-cell formation.^24^

An effective RSV vaccine will preferably induce robust humoral (VNA titers [VNT]) and cellular (CD4+ and CD8+ T-cell) responses.^25, 26^ Subunit vaccines based on pre-fusion conformation-stabilized RSV F protein (RSV preF protein) have shown robust induction of RSV VNT in preclinical models,^27, 28^ and phase 1 and 2 clinical trials have demonstrated their immunogenicity and tolerability.^29, 30^ Ad26.RSV.preF is a recombinant, replication-incompetent, adenovirus type 26 (Ad26)–vector encoding a conformation-stabilized RSV preF protein that effectively induces both humoral and cellular responses in animal models.^15, 27^ Ad26.RSV.preF has been evaluated for safety and immunogenicity in clinical trials; a single dose of Ad26.RSV.preF was safe and induced lasting immune responses in older adults (age ≥60 years)^31^ and demonstrated protection against RSV in a recent human challenge study of adults (ages 18-50 years).^32^

The combined data from nonclinical and clinical studies indicate that while Ad26.RSV.preF induces robust humoral and cellular immune responses, RSV preF protein subunit vaccines appear to induce superior humoral immune responses compared with Ad26.RSV.preF.^27^ To date, no published work has investigated the advantages of combining an RSV protein subunit vaccine with a vector-based vaccine. Here, we assessed the immunogenicity and efficacy of a co-administered combination of Ad26.RSV.preF and RSV preF protein and compared these with the responses to the individual components.

## Results

### RSV preF protein significantly contributes to humoral immune responses, whereas Ad26.RSV.preF strongly induces cellular responses when administered as a combination in naïve mice

To evaluate the potential advantages of combining Ad26.RSV.preF and RSV preF protein in a single vaccine, different dose levels of RSV preF protein (0.015 µg, 0.15 µg, 1.5 µg, and 15 µg) were combined with a low dose (1×10^8^ viral particles [vp]) of Ad26.RSV.preF and immunogenicity was compared with Ad26.RSV.preF alone in RSV-naïve mice. Additionally, a combination of high-dose Ad26.RSV.preF (1×10^9^ vp) and high-dose RSV preF protein (15 µg) was evaluated. Prime-boost immunization with combination regimens at Weeks 0 and 4 induced strong humoral and cellular immune responses measured at Week 6 (**Figure 1**). Mice immunized with Ad26.RSV.preF/RSV preF protein combinations showed significantly higher VNT against RSV CL57 compared with mice receiving Ad26.RSV.preF alone (P ≤ 0.02 for all RSV preF protein doses), with no clear difference in VNT between protein doses that ranged from 0.015 µg to 15 µg (**Figure 1a**). Similar findings were observed for binding antibodies against RSV preF protein measured by enzyme-linked immunosorbent assay (ELISA; **Figure 1b**). In addition, the Ad26.RSV.preF (1×10^9^ vp)/RSV preF protein (15 µg) combination vaccine induced significantly higher VNT and RSV preF binding antibodies compared with the Ad26.RSV.preF (1×10^8^ vp)/RSV preF protein (15 µg) combination vaccine (P = 0.019 and P < 0.001, respectively), demonstrating the dose-dependent contribution of Ad26.RSV.preF to vaccine-induced humoral responses.

**Figure 1.**
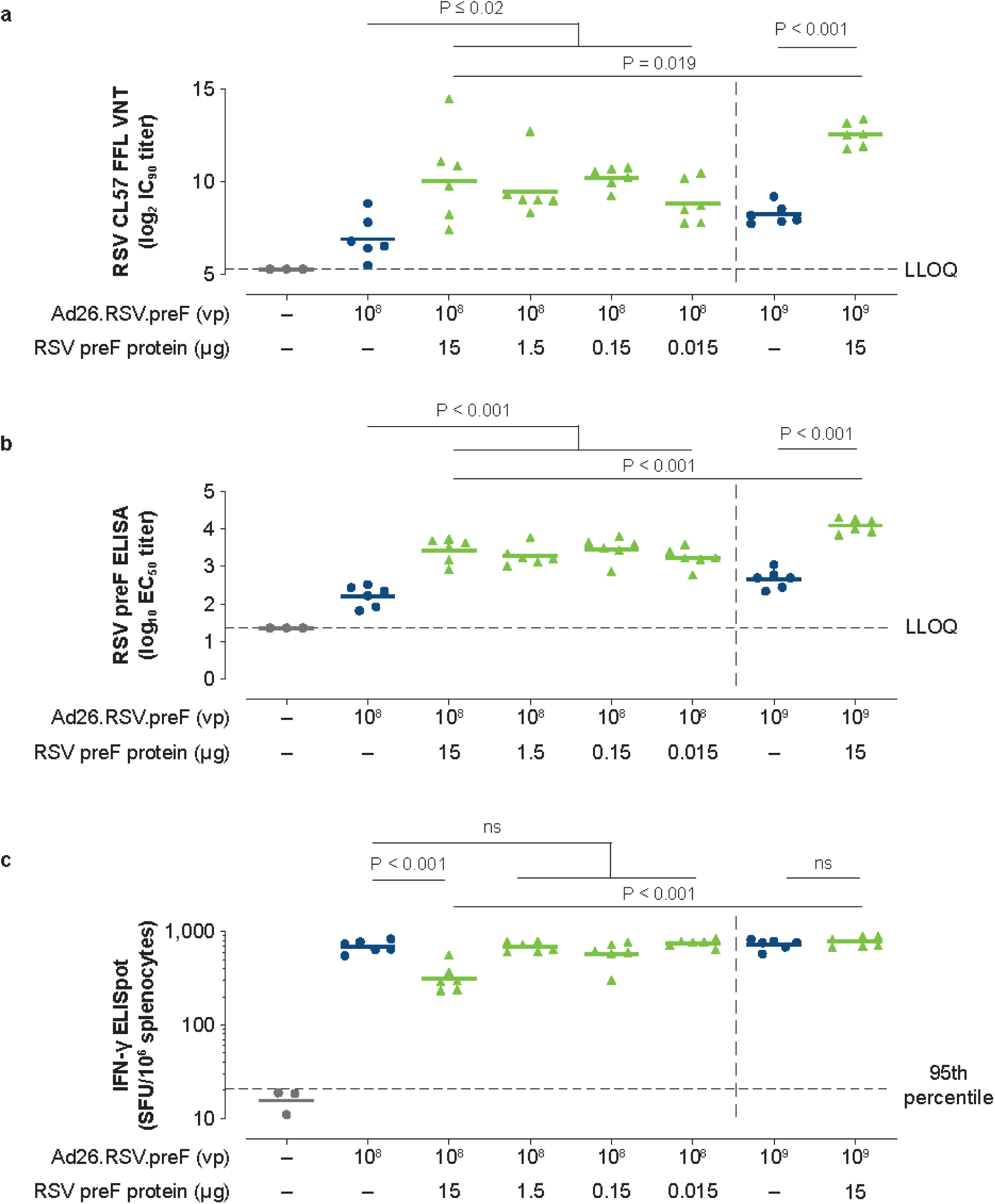
Humoral and cellular immunogenicity in naïve mice. Naïve female BALB/c mice were immunized at Week 0 and Week 4 with Ad26.RSV.preF (1×10^8^ or 1×10^9^ vp), Ad26.RSV.preF (1×10^8^ vp) combined with RSV preF protein (15, 1.5, 0.15 or 0.015 µg), or Ad26.RSV.preF (1×10^9^ vp) combined with RSV preF protein (15 µg; n = 6 each group). VNT against RSV CL57, expressed as log_2_ IC_90_ titers, using an FFL virus neutralization assay **(a)** and RSV preF–binding antibody titers, expressed as log_10_ EC_50_ titers **(b)**, were measured in serum at Week 6. Horizontal bars indicate mean titers per group, and dotted lines indicate the LLOQ or the LOD. Cellular responses were measured by IFN-γ ELISpot in splenocytes at Week 6 (**c**), and the responses are shown as SFU per million splenocytes. The horizontal lines indicate the geometric mean titer per group, and the dotted line indicates the 95^th^ percentile of the response of unstimulated cells. Ad26, adenovirus type 26; EC_50_, 50% effective concentration; ELISA, enzyme-linked immunosorbent assay; ELISpot, enzyme-linked immune absorbent spot; FFL, firefly luciferase; IC_90_, 90% inhibitory concentration; IFN-γ, interferon-γ; LLOQ, lower limit of quantification; LOD, limit of detection; ns, not significant; preF, prefusion conformation–stabilized RSV F protein; RSV, respiratory syncytial virus; SFU, spot-forming units; VNT, virus neutralization titers; vp, viral particles.

Prime-boost immunization with Ad26.RSV.preF alone induced robust cellular immune responses, as measured in splenocytes by RSV F–specific interferon-γ (IFN-γ) enzyme-linked immune absorbent spot (ELISpot; **Figure 1c**). There was no significant difference in cellular responses induced by low-dose Ad26.RSV.preF (1×10^8^ vp) alone or low-dose Ad26.RSV.preF combined with RSV preF protein at dose levels ranging from 0.015 µg to 1.5 µg. High-dose (15 µg) RSV preF protein combined with low-dose Ad26.RSV.preF showed a significantly lower cellular response compared with Ad26.RSV.preF alone. This was not observed with high-dose Ad26.RSV.preF (1×10^9^ vp), suggesting a dose-dependent effect of the adenoviral vector and the existence of an Ad26.RSV.preF/RSV preF protein dosage eliciting an optimal immune response. Further, the Ad26.RSV.preF (1×10^9^ vp)/RSV preF protein (15 µg) combination vaccine induced a significantly greater cellular immune response compared with the Ad26.RSV.preF (1×10^8^ vp)/RSV preF protein (15 µg) combination vaccine (P < 0.001), demonstrating the importance of Ad26.RSV.preF to vaccine-induced cellular immune responses.

### Ad26.RSV.preF/RSV preF protein combinations induce antibodies that effectively engage Fcγ receptors in an antibody-dependent cellular cytotoxicity reporter assay

RSV-naïve mice were immunized with 3 different dose levels of Ad26.RSV.preF alone (1×10^8^ vp, 1×10^9^ vp, and 1×10^10^ vp), RSV preF protein alone (0.15 µg, 1.5 µg, and 15 µg), and Ad26.RSV.preF/RSV preF protein combinations at Weeks 0 and 6. At Week 8, dose-dependent IgG1 and IgG2a antibody responses were observed for all vaccine regimens evaluated. In agreement with prior work,^27^ Ad26.RSV.preF alone induced high levels of IgG2a antibodies, whereas RSV preF protein alone induced an IgG1-biased response (**Figures 2a** and **2b**). Induction of both IgG subclasses was maintained by Ad26.RSV.preF/RSV preF protein combinations. In an antibody-dependent cellular cytotoxicity (ADCC) reporter assay using FcγRIV-transfected Jurkat cells and RSV-infected A549 cells as target cells, pooled serum from mice immunized with Ad26.RSV.preF and Ad26.RSV.preF/RSV preF protein combinations showed robust FcγRIV activation (**Figure 2c**). In contrast, pooled serum from mice immunized with RSV preF protein did not elicit any measurable response in the assay. Antibodies elicited by Ad26.RSV.preF alone, RSV preF protein alone, and the Ad26.RSV.preF/RSV preF protein combination demonstrated robust binding to RSV-infected A549 cells (**Figure 2d**). These findings correspond with vaccine-induced IgG1 and IgG2a antibody levels and demonstrate that the Ad26.RSV.preF/RSV preF protein combination maintains the robust Fc-mediated effector functions induced by the Ad26.RSV.preF component.

**Figure 2.**
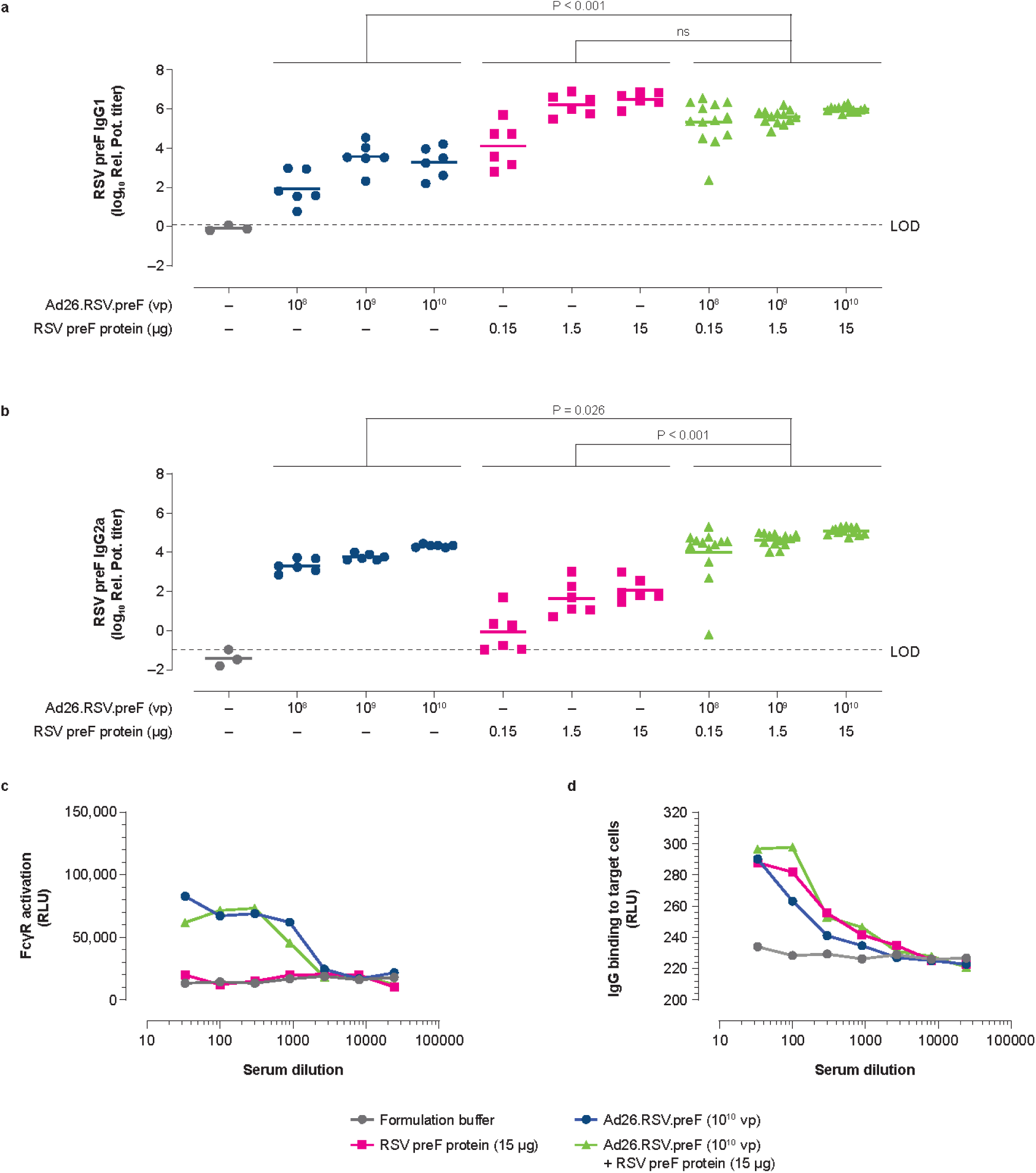
IgG subclass responses and FcγR engagement of antibodies induced by Ad26.RSV.preF/RSV preF protein combinations in naïve mice. Naïve female BALB/c mice were immunized at Week 0 and Week 6 with Ad26.RSV.preF (1×10^8^, 1×10^9^, or 1×10^10^ vp; n = 6 in each group), RSV preF protein (0.15, 1.5, or 15 µg; n = 6 each), or Ad26.RSV.preF combined with RSV preF protein at varying dose levels (n = 13 each). IgG1 (**a**) and IgG2a (**b**) preF-binding antibody titers, expressed as relative potency titer, were measured in serum by ELISA at Week 8. Horizontal lines indicate mean antibody responses per group, and the dotted line indicates the LOD, specified as the titer of the sample with the highest titer in the formulation buffer control group. Serum obtained at Week 8 from mice immunized with the highest vaccine concentration (Ad26.RSV.preF [1×10^10^ vp] and RSV preF protein [15 ug], alone or in combination) was pooled and used to measure FcγRIV activation in an ADCC reporter assay (**c**), expressed as RLU. Total IgG binding to RSV F–expressing target cells (**d**) was measured by luminescence using AF488-conjugated goat anti-mouse IgG. Ad26, adenovirus type 26; ADCC, antibody-dependent cellular cytotoxicity; ELISA, enzyme-linked immunosorbent assay; FcγR, Fcγ receptor; FcγRIV, Fcγ receptor IV; IgG, immunoglobulin G; LOD, limit of detection; ns, not significant; preF, prefusion conformation– stabilized RSV F protein; Rel. Pot., relative potency; RLU, relative light units; RSV, respiratory syncytial virus; vp, viral particles.

### Ad26.RSV.preF/RSV preF protein combinations induce humoral and cellular immune responses in RSV pre-exposed mice

To mimic the RSV pre-exposed status of most adult humans, mice were pre-exposed intranasally (i.n.) with RSV A2 (5×10^5^ plaque-forming units [pfu]) at 6 to 8 weeks of age and received a single immunization with Ad26.RSV.preF (1×10^8^ vp or 1×10^9^ vp), RSV preF protein (0.15 µg), or Ad26.RSV.preF/RSV preF protein combinations at 18 months of age (Week 0). Immunization with Ad26.RSV.preF/RSV preF protein combinations induced robust VNT responses in RSV pre-exposed mice at Week 5, with significantly higher responses compared with Ad26.RSV.preF alone (**Figure 3a**). Similar findings were observed for RSV preF–binding IgG antibodies (**Figure 3b**). Ad26.RSV.preF administered alone induced robust cellular immune responses, as measured by RSV F–specific IFN-γ ELISpot (**Figure 3c**) and intracellular cytokine staining (**Supplementary Figure 1**); the addition of RSV preF protein to Ad26.RSV.preF at the evaluated doses did not impact the cellular immune responses induced by Ad26.RSV.preF alone. Similar to observations in naïve mice, the contribution of the RSV preF protein to cellular responses was negligible.

**Figure 3.**
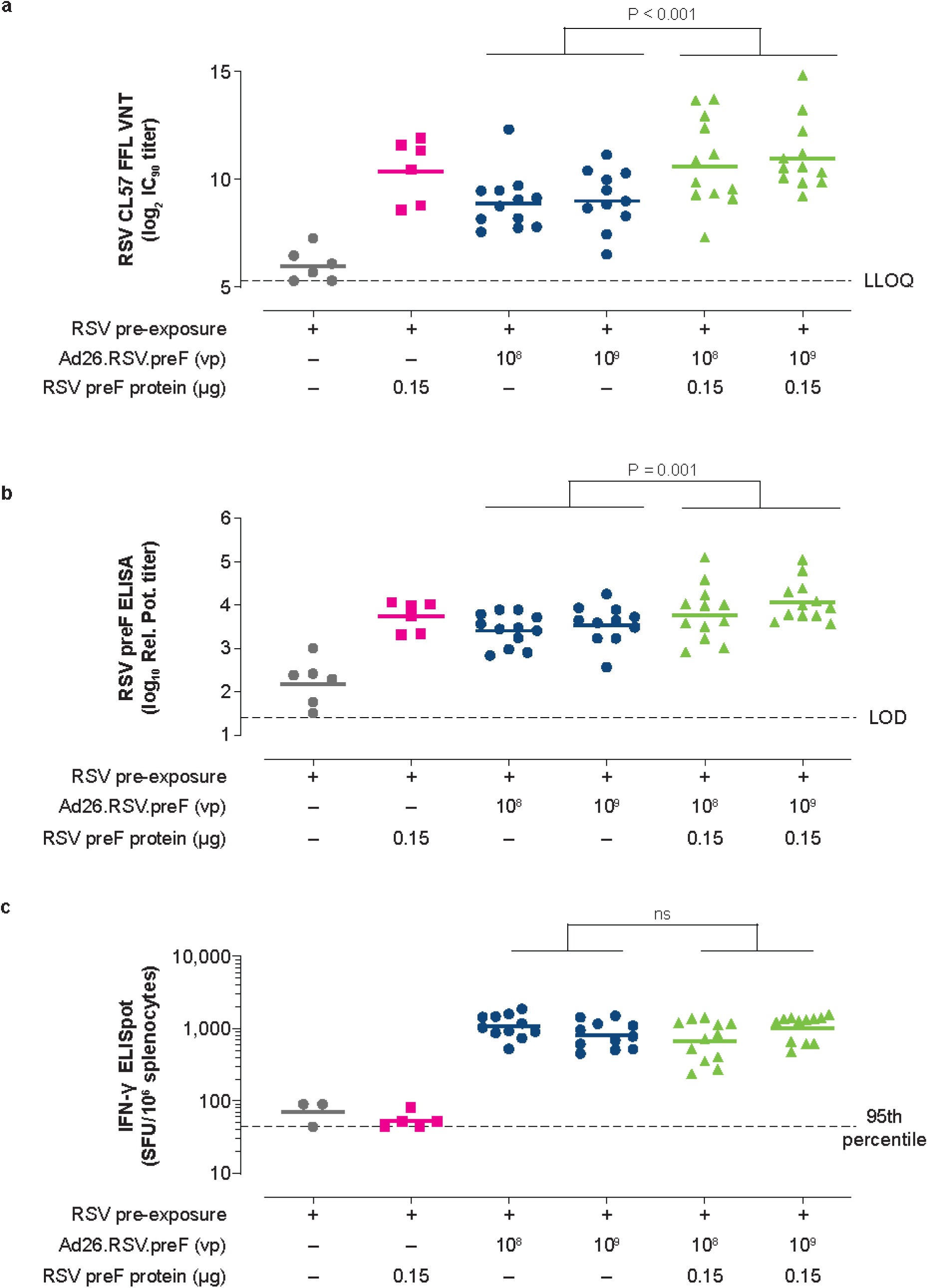
Humoral and cellular immunogenicity in RSV pre-exposed mice. Naïve female BALB/c mice were intranasally pre-exposed with RSV A2 (5×10^5^ pfu) at Week –79 and immunized intramuscularly at Week 0 with Ad26.RSV.preF (1×10^8^ or 1×10^9^ vp; n = 12 in each group), RSV preF protein (0.15 µg; n = 6), or Ad26.RSV.preF (1×10^8^ or 1×10^9^ vp) combined with RSV preF protein (0.15 µg; n = 12 each group). Negative control animals received formulation buffer (n = 6). VNT against RSV CL57, expressed as log_2_ IC_90_ titers, were measured in serum at Week 5 **(a)**. Horizontal bars indicate mean VNT per group, and the dotted line indicates the LLOQ. RSV preF total IgG-binding antibody titers (**b**), expressed as log_10_ relative potency titers, were also measured in serum at Week 5. Horizontal bars indicate mean titers per group, and the dotted line indicates the LOD. Cellular responses were measured by IFN-γ ELISpot in splenocytes at Week 5 (**c**), and the responses are expressed as SFU per million splenocytes. The horizontal lines indicate the geometric mean response per group, and the dotted line indicates the 95^th^ percentile of the response of unstimulated cells. Ad26, adenovirus type 26; ELISA, enzyme-linked immunosorbent assay; ELISpot, enzyme-linked immune absorbent spot; FFL, firefly luciferase; IC_90_, 90% inhibitory concentration; IFN-γ, interferon-γ; IgG, immunoglobulin G; LLOQ, lower limit of quantification; LOD, limit of detection; ns, not significant; pfu, plaque-forming units; preF, prefusion conformation–stabilized RSV F protein; Rel. Pot., relative potency; RSV, respiratory syncytial virus; SFU, spot-forming units; VNT, virus neutralization titers; vp, viral particles.

These data demonstrate that Ad26.RSV.preF/RSV preF protein combinations elicit robust humoral and cellular responses in RSV pre-exposed mice, leveraging the advantages of the individual vaccine components.

### Protective efficacy of Ad26.RSV.preF/RSV preF protein combinations in cotton rats

To study the protective efficacy of the Ad26.RSV.preF/RSV preF protein combination, a naïve cotton rat challenge model was used. As RSV pre-exposure already effectively protects cotton rats against RSV challenge, vaccine efficacy could not be evaluated under pre-immune conditions in this model.^33^ Naïve cotton rats were immunized with subprotective doses of Ad26.RSV.preF (1×10^5^ vp and 1×10^6^ vp) and RSV preF protein (0.5 µg), alone and in combination, to evaluate the contributions of the individual components. Animals were challenged i.n. with RSV A2 at Day 49 and sacrificed at Day 54 for collection of lung tissue and nasal turbinates.

Cotton rats immunized with a low dose (1×10^5^ vp) of Ad26.RSV.preF alone were poorly protected against RSV in nasal and lung tissue, with 0/12 and 2/12 animals showing complete protection, respectively (**Figures 4a** and **4b**). This low protection corresponded with negligible VNT measured at Day 49 in prechallenge serum (**Figure 4c**). A higher dose (1×10^6^ vp) of Ad26.RSV.preF induced VNT and provided complete protection in the lungs in all animals; however, all animals demonstrated breakthrough infections in nasal tissue, but with lower viral loads compared with the mock-immunized control group. Animals receiving RSV preF protein (0.5 µg) alone did not show VNT above background levels and subsequently demonstrated only marginal protection, with high viral titers in both nasal and lung tissue. Immunization with Ad26.RSV.preF/RSV preF protein combinations resulted in significantly lower nasal viral loads compared with Ad26.RSV.preF alone, with 6/12 animals demonstrating complete protection in the Ad26.RSV.preF 1×10^6^ vp/RSV preF protein 0.5-µg group. However, this difference in protection was not reflected in vaccine-elicited VNT, as there were no significant differences in VNT between animals immunized with Ad26.RSV.preF alone and those immunized with Ad26.RSV.preF/RSV preF protein combinations (**Figure 4c**).

**Figure 4.**
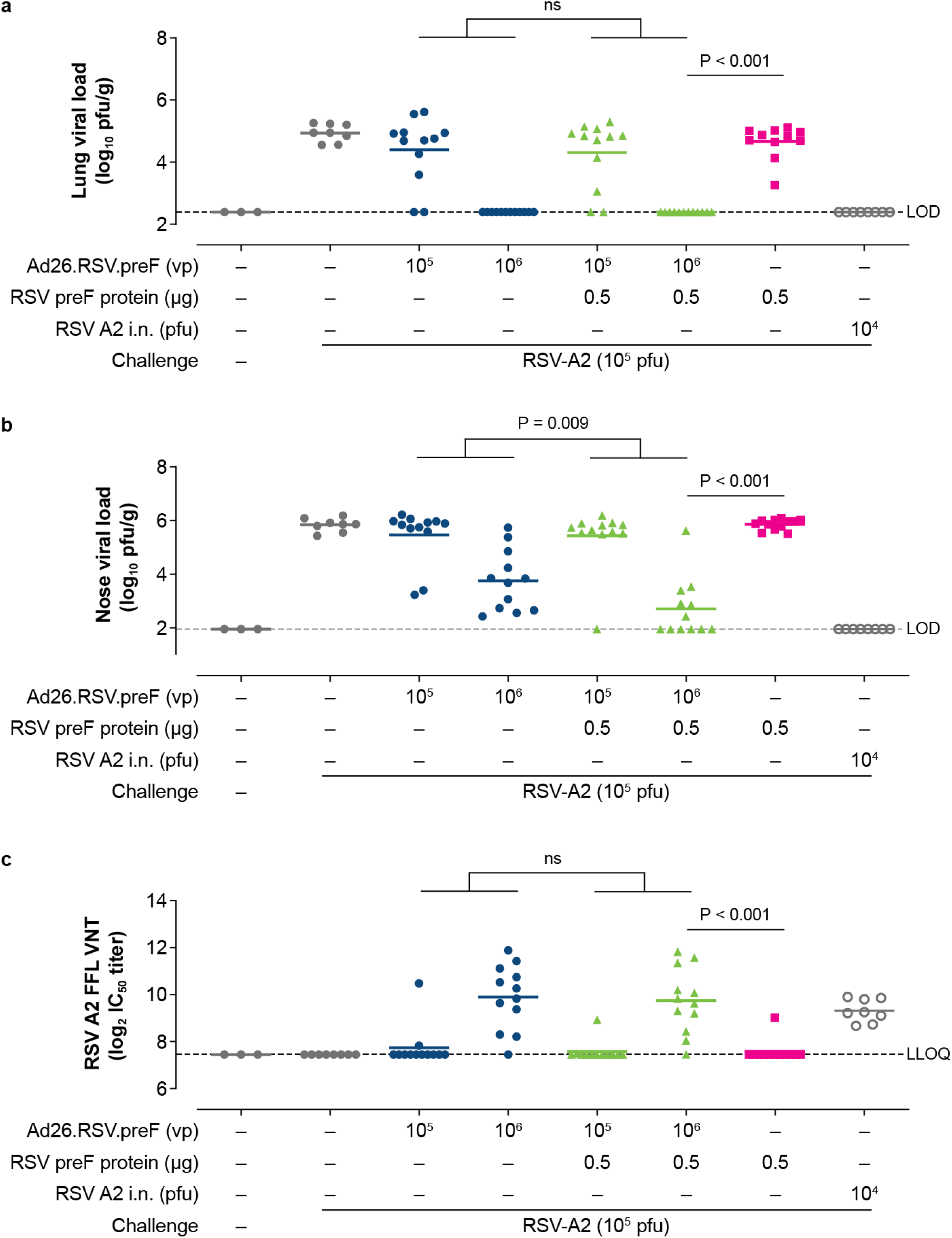
Immunogenicity and protective efficacy in cotton rats. Naïve female cotton rats were immunized intramuscularly at Week 0 with Ad26.RSV.preF (1×10^5^ or 1×10^6^ vp; n = 12 in each group), RSV preF protein (0.5 µg; n = 12), or Ad26.RSV.preF (1×10^5^ or 1×10^6^ vp) combined with RSV preF protein (0.5 µg; n = 13 in each group). Negative control groups received phosphate buffered saline (n = 3 and n = 8, assay control and challenge control, respectively) and the positive control group received i.n. RSV A2 (1×10^4^ pfu; n = 8). Cotton rats were challenged i.n. with RSV A2 (1×10^5^ pfu) at Day 49 and Day 54. Viral load was measured in lung (**a**) and nose homogenates (**b**) and expressed as log_10_ pfu/g tissue. Horizontal lines indicate the mean viral load per group, and the dotted line indicates the LOD. RSV A2 VNT, expressed as log_2_ IC_50_, were measured in prechallenge serum samples obtained at Day 49 (**c**). Horizontal lines indicate mean VNT per group, and the dotted line indicates the LLOQ. Ad26, adenovirus type 26; FFL, firefly luciferase; IC_50_, half-maximum inhibitory concentration; i.n., intranasally; LLOQ, lower limit of quantification; LOD, limit of detection; ns, not significant; pfu, plaque-forming units; preF, prefusion conformation–stabilized RSV F protein; RSV, respiratory syncytial virus; VNT, virus neutralization titers; vp, viral particles.

There was a significant reduction in nasal and lung viral loads in animals immunized with the Ad26.RSV.preF (1×10^6^ vp)/RSV preF protein (0.5 µg) combination compared with RSV preF protein alone (P <0.001 for both nasal and lung viral titers), and the former group demonstrated significantly higher VNT (P <0.001).

### Immunization with Ad26.RSV.preF/RSV preF protein combinations shows robust and durable immune responses in RSV pre-exposed African green monkeys

Ad26.RSV.preF/RSV preF protein combinations were evaluated in RSV pre-exposed adult and aged African green monkeys (AGMs), and immune responses were compared with responses induced by the individual vaccine components. AGMs were pre-exposed i.n. with RSV Memphis 37 (7.5×10^5^ pfu) 19 weeks prior to immunization. Inoculation elicited low levels of VNT against RSV CL57 at 2, 14, and 18 weeks postinfection, confirming successful i.n. exposure. Nineteen weeks postinfection, animals were immunized with Ad26.RSV.preF (1×10^11^ vp) alone, RSV preF protein (150 µg) alone, and combinations comprising Ad26.RSV.preF (1×10^11^ vp) with varying doses of RSV preF protein (15 µg, 50 µg, and 150 µg).

Ad26.RSV.preF, RSV preF protein, and combinations all induced robust VNT against RSV CL57 after 1 immunization, with all groups showing a peak response at 2 weeks postimmunization (**Figure 5a**). The geometric mean fold rise (GMFR) in VNT, measured in individual animals at their peak response at Week 2 or 4, was 77.0, 113.2, and 97.6 for Ad26.RSV.preF, RSV preF protein, and all 3 Ad26.RSV.preF/RSV preF protein combinations combined, respectively (at Week 4 the GMFR was 55.2, 43.1, and 56.0, respectively). No significant differences in RSV CL57 VNT were observed between the group receiving Ad26.RSV.preF only and groups receiving Ad26.RSV.preF/RSV preF protein combinations at any time point tested prior to boost immunization at Week 60. Although animals receiving RSV preF protein (150 µg) alone showed similar peak responses, VNT were significantly lower compared with the Ad26.RSV.preF groups after Week 4, suggesting reduced durability of the response in this group.

**Figure 5.**
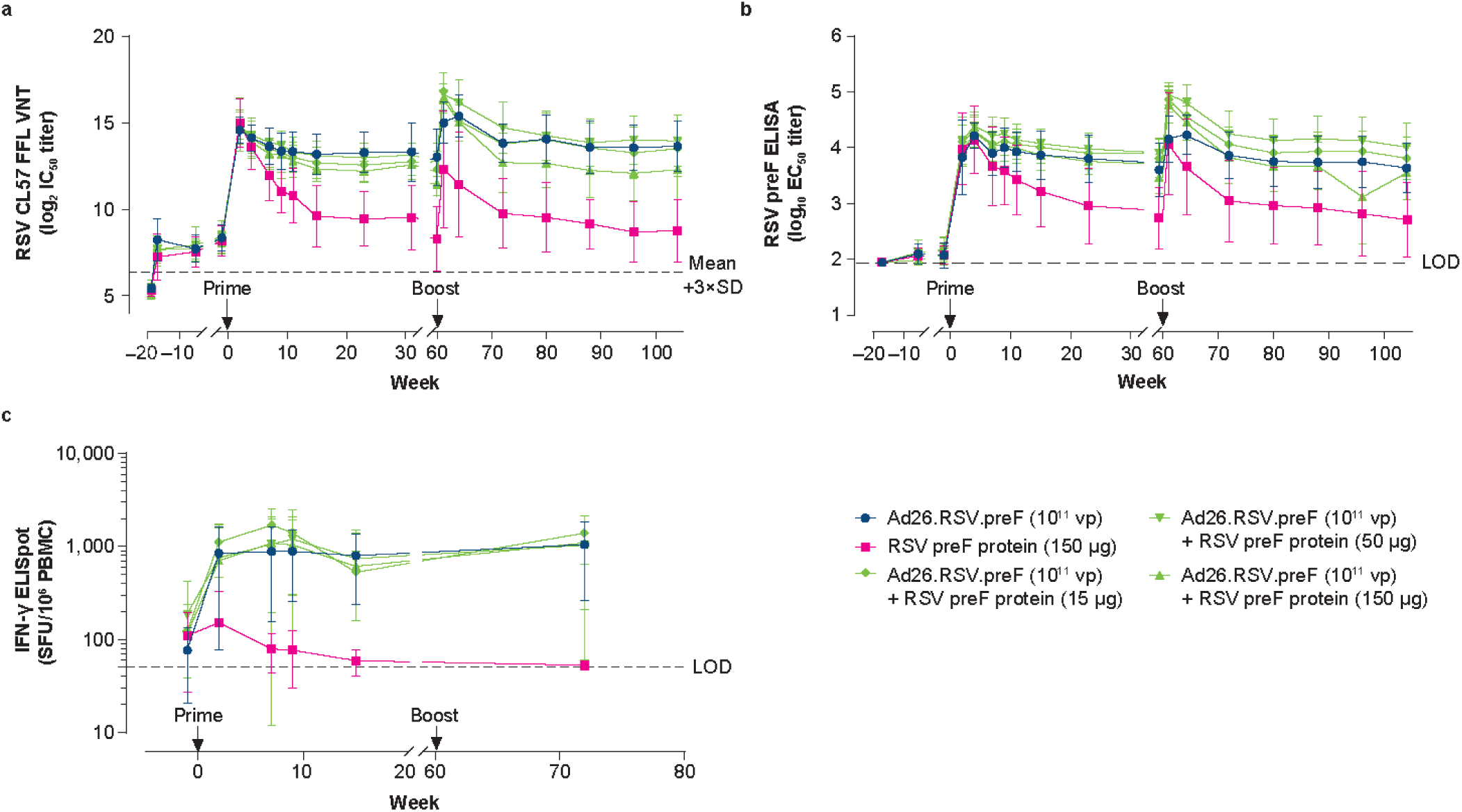
Humoral and cellular immunogenicity measured in African green monkey serum. Female African green monkeys were pre-exposed i.n. with RSV Memphis 37 (7.5×10^5^ pfu) at Week –19 and immunized intramuscularly with Ad26.RSV.preF (1×10^11^ vp; n = 11); RSV preF protein (150 µg; n = 4); or a combination of Ad26.RSV.preF (1×10^11^ vp) with RSV preF protein (15, 50, or 150 µg; n = 7 in each group) at Week 0 with homologous boost immunization at Week 60. RSV CL57 VNT, expressed as log_2_ IC_50_, were measured in serum samples; dots represent mean VNT per group; error bars indicate SD (**a**). LOD was determined as 3×SD above mean VNT of Week –19 serum samples and indicated by a dotted line. RSV preF IgG-binding titers, expressed as log_10_ EC_50_, were measured by ELISA in serum samples (**b**). Dots represent mean titers per group; error bars indicate SD. LOD was determined as log_10_ of the inverse of the lowest serum dilution. PBMCs were obtained before immunization (but after pre-exposure) and after immunization at Weeks 2, 7, 9, 15, and 72 for measurements of cellular immune responses by IFN-γ ELISpot (**c**), expressed as SFU/10^6^ PBMCs. Each dot represents geometric mean responses per group, and error bars indicate 95% CIs. LOD was set at 50 SFU/10^6^ PBMCs. Measurement of the animal showing the highest SFU counts at Weeks 2, 7, and 9 (in the group receiving Ad26.RSV.preF/RSV preF protein [15 μg]) is missing at Week 15 due to too many spots. Ad26, adenovirus type 26; CI, confidence interval; EC_50_, 50% effective concentration; ELISA, enzyme-linked immunosorbent assay; ELISpot, enzyme-linked immune absorbent spot; FFL, firefly luciferase; IC_50_, half-maximum inhibitory concentration; IFN-γ, interferon-γ; IgG, immunoglobulin G; i.n., intranasally; LOD, limit of detection; PBMC, peripheral blood mononuclear cell; pfu, plaque-forming units; preF, prefusion conformation–stabilized RSV F protein; RSV, respiratory syncytial virus; SD, standard deviation; SFU, spot-forming units; VNT, virus neutralization titers; vp, viral particles.

AGMs immunized with Ad26.RSV.preF and the combination vaccine showed robust VNT responses that were stable up to Week 60. Homologous boost immunization at Week 60 induced a significant boost response in groups receiving Ad26.RSV.preF alone and Ad26.RSV.preF/RSV preF protein combinations (comparing Week 60 and Week 61; P ≤0.035 for all; **Figure 5a**). RSV preF protein alone also induced a boost response of 17-fold; however, this response did not reach statistical significance (P = 0.061) due to the low number of animals (n = 4) and high variation in this group. Prior to boost immunization, RSV preF protein showed significantly lower VNT compared with Ad26.RSV.preF alone and the Ad26.RSV.preF (1×10^11^ vp)/RSV preF protein (150 µg) combination (P <0.001 and P = 0.019, respectively); this difference remained significant at Week 61 after homologous boost immunization (P = 0.02 and P = 0.001, respectively; **Figure 5a**). The GMFR to peak VNT responses, measured either at Week 61 or 64, was 5.9 and 22.0 for Ad26.RSV.preF alone and all 3 Ad26.RSV.preF/RSV preF protein combinations combined, respectively, suggesting a substantial contribution of the protein component to the VNT boost response. No significant differences in VNT responses were observed between groups receiving Ad26.RSV.preF alone and Ad26.RSV.preF/RSV preF protein combinations at later time points (Week 64 to Week 104), whereas the RSV preF protein alone group showed reduced VNT responses. Observed vaccine-induced RSV preF–binding IgG titers largely matched those for RSV VNT (**Figure 5b**).

Prior to immunization, all animals exhibited low RSV F–specific IFN-γ ELISpot responses resulting from the i.n. exposure at Week –19. Immunization with Ad26.RSV.preF alone and Ad26.RSV.preF/RSV preF protein combinations, but not RSV preF protein alone, elicited robust cellular responses at Weeks 2, 7, 9, and 15. These robust cellular responses were maintained at Week 72 (ie, 12 weeks after boost immunization) in animals who received Ad26.RSV.preF alone and Ad26.RSV.preF/RSV preF protein combinations (**Figure 5c**). ICS analysis showed RSV F– specific CD4+ and CD8+ T cells in animals before immunization (Week –1), likely as a result of RSV pre-exposure at Week –19. Immunization with both Ad26.RSV.preF alone and Ad26.RSV.preF/RSV preF protein combinations induced CD8+ T cells expressing IFN-γ, interleukin (IL)-2, and tumor necrosis factor-(TNF-) α, with high variation within groups (no statistical analysis performed; **Supplementary Figure 2**). RSV preF protein alone showed minimal induction of CD8+ T cells, with marginal induction of CD8+ T cells expressing IL-2. Ad26.RSV.preF/RSV preF protein combinations showed marginal induction of CD4+ T-cell responses, with no apparent differences between groups, whereas RSV preF protein alone induced no CD4+ T cell response above baseline (**Supplementary Figure 3**).

RSV preF IgA and IgG titers were measured in bronchoalveolar lavage (BAL) samples gathered from AGMs before and 2 weeks after prime immunization (no BAL samples were obtained after boost immunization at Week 60). In addition, RSV preF IgA titers were measured in nasal swabs preimmunization and at Weeks 2, 60, 61, and 104. Two weeks after prime immunization, mean RSV preF IgA and IgG titers in the BAL and preF IgA titers in nasal swabs were increased in all vaccinated groups compared with baseline, and no differences were observed between groups (**Figure 6**). From Week 2 to Week 60, there was a reduction in IgA titers in nasal swabs in all groups, although titers generally remained above baseline. Boost immunization at Week 60 increased RSV preF nasal swab titers in all groups, although no statistical comparisons were made due to low sample sizes resulting from the exclusion of blood-contaminated samples. However, all groups showed elevated nasal swab IgA titers through the end of the study (Week 104), with no clear differences between groups.

**Figure 6.**
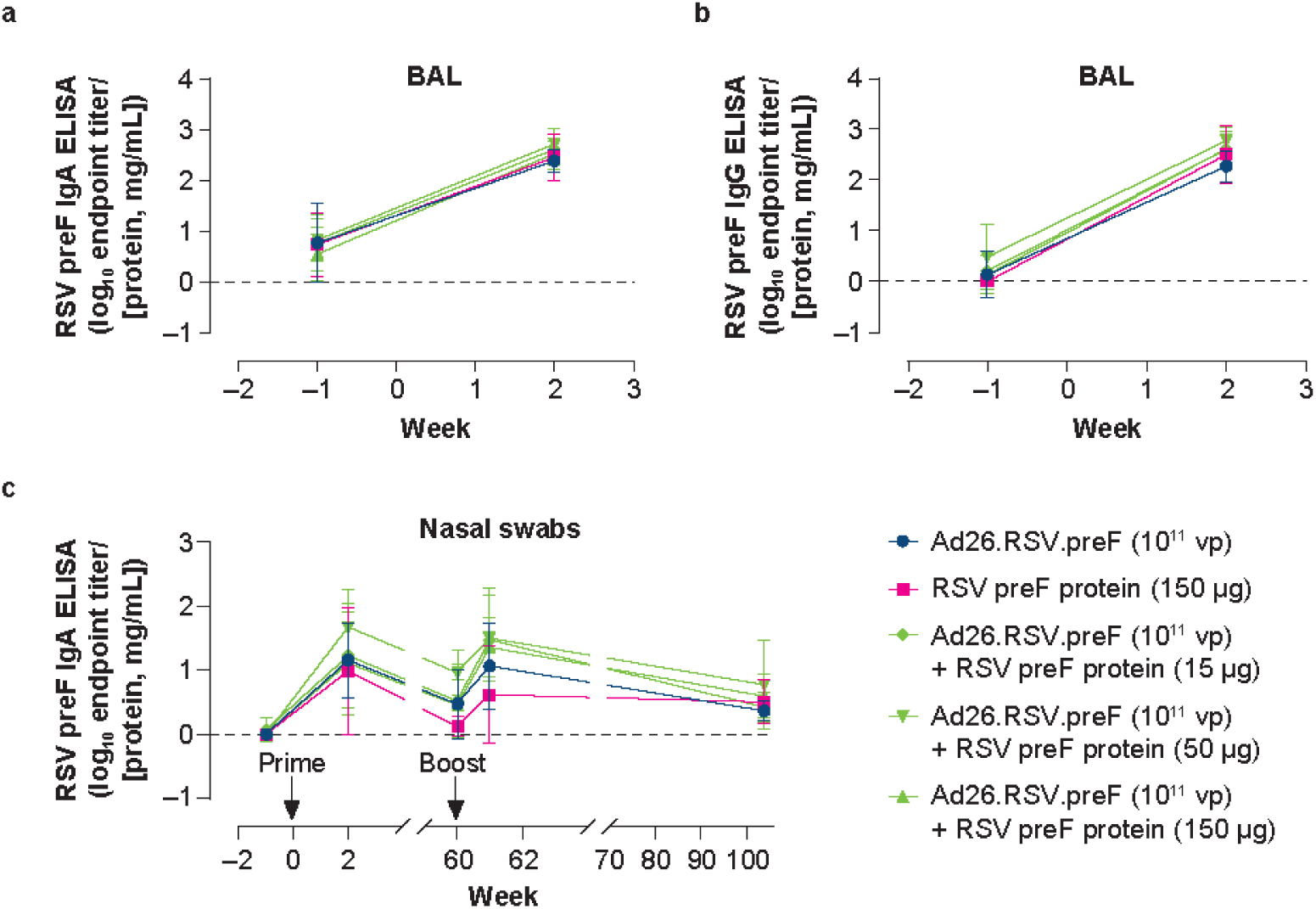
RSV preF IgA and IgG titers measured in nasal swabs and BAL of African green monkeys. African green monkeys were immunized as described in Figure 5. RSV preF IgA **(a)** and IgG **(b)** antibody-binding titers, expressed as endpoint titer (ELISA) divided by total protein concentration measured at OD280, in BAL samples obtained before immunization and after immunization at Week 2. No BAL samples were collected after boost immunization. RSV preF IgA ELISA titers (**c**) were measured in nasal swabs obtained before immunization and at Weeks 2, 60, 61, and 104. Ad26, adenovirus type 26; BAL, bronchoalveolar lavage; ELISA, enzyme-linked immunosorbent assay; IgA, immunoglobulin A; IgG, immunoglobulin G; preF, prefusion conformation–stabilized RSV F protein; RSV, respiratory syncytial virus; vp, viral particles.

## Discussion

We have demonstrated that immunization with a combination of an adenoviral vector (Ad26.RSV.preF) and RSV preF protein induces robust humoral and cellular immune responses, both in RSV pre-exposed mice and AGMs. In AGMs, the antibody responses showed high durability up to 60 weeks and strong cellular immune responses were elicited. Immunologic analysis in mice and AGMs showed that antibody durability, Fc-mediated effector functions, and cellular immune responses were primarily attributable to the Ad26.RSV.preF component, while the RSV preF protein contributed to robust antibody induction. These data confirm prior studies with each vaccine component^27^ and highlight the advantages of a combination vaccine for robust induction of durable humoral and cellular immune responses.

Immune CoP for RSV infection remain ill-defined because RSV induces only modest and transient immunity and people remain susceptible to infection and RSV disease throughout life.^34^ Human challenge models have provided evidence that both humoral and cellular immune responses, particularly in the mucosal compartment, play an important role in protection against RSV infection. RSV-specific nasal IgA antibodies at the time of challenge have been shown to correlate with protection against infection,^19^ and CD8+ T cells in the airway have been associated with lower symptom severity and reduced viral load after subsequent challenge.^21^ Preclinical studies also support the role of both humoral and cellular immune responses for preventing RSV infection and reducing RSV disease severity.^22, 25, 35–37^ In mouse models, T cells effectively clear RSV infection from the lungs, and depletion of both CD4+ and CD8+ T cells has been shown to prolong RSV infection after challenge.^22, 35^ In a human challenge study, immunization with Ad26.RSV.preF resulted in reduced viral load, RSV infections, and disease severity after challenge with RSV Memphis 37b; additionally, increased serum RSV A2 VNT was associated with reduced probability of infection.^32^ A recent study investigating CoPs in AGMs across 6 different investigational RSV vaccines found that RSV preF–specific mucosal IgA antibodies were correlated with protection from both upper and lower respiratory RSV infections; VNAs and ADCC correlated with viral control in the upper respiratory tract, and IgG-mediated Fc-effector functions were shown to be important for RSV reduction in the lower respiratory tract.^36^ Collectively, both human and animal studies demonstrated that different components of the immune system (ie, humoral and cellular responses in both the mucosal and systemic compartments) contributed to effective control of RSV replication. In the current study, we demonstrated that the Ad26.RSV.preF/RSV preF protein combination vaccine elicits broadly effective immune responses, which is especially important for an RSV vaccine to maintain efficacy in heterogeneous adult and older adult populations.

Because nearly every person experiences RSV infection by 2 years of age and re-infection frequently occurs throughout life,^38^ we used aged (18 month old), RSV pre-exposed mice to evaluate the immunogenicity of the Ad26.RSV.preF/RSV preF protein combination. Similar to older human adults, mice ≥18 months of age show diminished immunity, with impacts of immune senescence on both the humoral and cellular compartments of the adaptive immune system.^33, 39^ For a vaccine targeting older adults, it is important to demonstrate robust vaccine-induced immune responses in a suitable animal model. In this study, the addition of RSV preF protein to Ad26.RSV.preF increased vaccine-induced humoral immune responses in aged, RSV pre-exposed mice; further, Ad26.RSV.preF/RSV preF protein combinations induced robust cellular immune responses in these mice, similar to Ad26.RSV.preF alone. The data generated in this pre-exposure model highlight the benefits of the Ad26.RSV.preF/RSV preF protein combination, which induces robust immune responses in both the humoral and cellular immunity compartments.

In the RSV A2 cotton rat challenge model, the Ad26.RSV.preF/RSV preF protein combination vaccine reduced viral loads in both the lungs and the nose. In the nose, there was a significantly lower viral load in animals immunized with Ad26.RSV.preF/RSV preF protein combinations compared with Ad26.RSV.preF alone. This was, however, not associated with the level of VNT in prechallenge serum, suggesting that other immune parameters play a role. In RSV pre-exposed adult and aged AGMs, Ad26.RSV.preF/RSV preF protein combinations induced mucosal IgA antibodies both in BAL fluid and nasal swabs. Because RSV-specific IgA antibodies have been suggested to correlate with protection in humans, it is plausible that they play a role in limiting viral load in the nose in the cotton rat challenge model.

IgG-mediated Fc-effector functions are likely to contribute to virus control as well, especially in the lower respiratory tract..^36, 40^ IgG subclasses play differential roles in host defense; the IgG2a subclass has demonstrated the ability to potently interact with Fc receptors.^41^ FcγRIV, expressed by myelocytes, is an activating Fc receptor that exhibits intermediate binding affinity for IgG2a and IgG2b subclass antibodies and no binding affinity for IgG1.^41^ A preclinical study in BALB/c mice demonstrated that Ad26.RSV.preF elicits RSV F–specific IgG2a subclass antibodies that induce Fc-mediated effector functions through FcγRIV in an ADCC reporter assay; in contrast, RSV preF protein primarily elicits IgG1 subclass antibodies that do not mediate effector functions via FcγRIV.^27^ In this study, IgG subclasses induced by the individual components in naïve BALB/c mice were maintained by the combination vaccine, resulting in a robust, multifaceted immune response.

In the AGM study, there was a robust induction of VNT, with no significant differences between animals receiving Ad26.RSV.preF/RSV preF protein combinations or Ad26.RSV.preF alone. Fold-increases in VNT, measured at 4 weeks after immunization, were 55.2 and 56.0 for Ad26.RSV.preF alone and Ad26.RSV.preF/RSV preF protein combinations, respectively. In contrast, Ad26.RSV.preF and Ad26.RSV.preF/RSV preF protein combinations induced fold-increases in RSV A2 VNT of 2.8 and 10.3, respectively, after 4 weeks in humans receiving the same dose levels.^42^ Differences in VNT fold-increases between AGMs and humans might be due to the substantial difference in the pre-exposure history of both. While AGMs in this study received a single i.n. pre-exposure with RSV Memphis 37b that resulted in low but detectable antibody titers, humans have different and complex pre-exposure histories, likely with repeated re-infections with multiple different strains and overall higher antibody titers. Another discrepancy between AGM and human studies is that the clinical study showed the added benefit of RSV preF protein for optimal induction of a neutralizing-antibody response,^42^ whereas this was not apparent in the AGM study. Whether the difference in pre-exposure history is also underlying this discrepancy is currently unknown.

The robust induction of VNT in AGMs also showed high durability. VNT durability was attributed to the Ad26.RSV.preF component, as antibodies induced by immunization with RSV preF protein alone were substantially less durable. In a phase 1 clinical study, Ad26.RSV.preF was shown to elicit robust humoral and cellular immune responses that were durable for ≥2 years postvaccination.^31^ The effective induction of cellular immune responses by Ad26.RSV.preF likely contributed to the durability of humoral immune responses; RSV preF protein alone elicited minimal T-cell responses, and humoral durability was poor compared with Ad26.RSV.preF. Although nonadjuvanted RSV preF protein–based subunit vaccines have demonstrated robust humoral immune responses in clinical trials,^30, 43–45^ they generally do not induce effective T-cell responses. Adjuvants can improve cellular immunity elicited by subunit vaccines.

In the current study, Ad26.RSV.preF was the main contributor to cellular immune responses in both mice and AGMs. In contrast to subunit vaccines, vector-based vaccines are generally considered effective vaccine platforms for eliciting T-cell immunity.^46–50^ In mice, a relatively high concentration of RSV preF protein in the combination vaccine resulted in reduced T-cell responses compared with responses elicited by Ad26.RSV.preF alone, as measured by IFN-γ ELISpot and ICS. This reduction in cellular responses did not occur at relative RSV preF protein dose ratios equal to or less than that intended for human use (Ad26.RSV.preF [1×10^11^ vp]/ RSV preF protein [150 µg]). The reason for this effect is not clear; however, we hypothesize that high protein antigen concentrations may inhibit adenoviral vector transduction of target cells or create a microenvironment that affects antigen presentation. Overall, this may reduce induction of T-cell responses.

Here, we show for the first time that the addition of RSV preF protein to Ad26.RSV.preF improves the humoral responses associated with Ad26.RSV.preF, while preserving the robust cellular immune responses induced by Ad26.RSV.preF alone. Results were consistent between small-animal rodent models (ie, mice and cotton rats) and a nonhuman primate model, providing strong evidence for the potential efficacy of an Ad26.RSV.preF/RSV preF protein combination RSV vaccine for adults. Further studies evaluating the safety, immunogenicity, and efficacy of Ad26.RSV.preF/RSV preF protein combinations in humans are ongoing.

## Methods

### Experimental design

#### Immunogenicity in naïve and RSV pre-exposed mice

Experiments evaluating the immunogenicity of Ad26.RSV.preF, RSV preF protein, and combinations of both components in naïve mice were conducted as follows: BALB/c mice (Charles River Laboratories, Sulzveld, Germany; 6-8 weeks old; n = 6 per group; n = 3 in the control group) were immunized intramuscularly (i.m.) at Week 0 with Ad26.RSV.preF alone (1×10^8^ or 1×10^9^ vp), Ad26.RSV.preF/RSV preF protein (0.015 µg, 0.15 µg, 1.5 µg, or 15 µg) combinations, or phosphate-buffered saline (PBS) control, with a second homologous dose at Week 4. Serum samples (for VNT and ELISA measurements) and spleens (for RSV F–specific, IFN-γ ELISpot measurements) were collected at Week 6 for immunogenicity analyses.

Experiments evaluating the immunogenicity of Ad26.RSV.preF and RSV preF protein in RSV pre-exposed mice were conducted as follows: at Week –79 prior to immunization, female BALB/c mice (6-8 weeks old) were pre-exposed i.n. with 5×10^5^ pfu of RSV A2. At Week 0, RSV pre-exposed mice were immunized i.m. with Ad26.RSV.preF (1×10^8^ vp [n = 12] or 1×10^9^ vp [n = 12]), RSV preF protein (0.15 µg; n = 6), a combination of Ad26.RSV.preF/RSV preF protein (1×10^8^ vp/0.15 µg [n = 12] or 1×10^9^ vp/0.15 µg [n = 12]), or PBS (n = 5). Serum samples (for VNT and ELISA measurements) and spleens (for RSV F–specific, IFN-γ ELISpot measurements and ICS) were collected at Week 5 for immunogenicity analyses.

#### Immunogenicity and protective efficacy in cotton rats

Experiments evaluating the immunogenicity and protective efficacy of Ad26.RSV.preF and RSV preF protein in cotton rats were conducted as follows: female cotton rats (Sigmovir Biosystems, Inc., Rockville, MD, USA; 6-8 weeks old; n = 12 per group) were immunized i.m. at Day 0 with Ad26.RSV.preF alone (1×10^5^ or 1×10^6^ vp), RSV preF protein alone (0.5 µg), or a mix of Ad26.RSV.preF and RSV preF protein. Control groups received i.n. RSV A2 (1×10^4^ pfu) or i.n. PBS. Animals were challenged i.n. with RSV A2 (1×10^5^ pfu) at Day 49 and sacrificed at Day 54. Viral load was determined in lung and nose homogenates by plaque assay, and serum samples were collected at Days 0, 28, and 49. Cotton rat experiments were performed by Sigmovir Biosystems, Inc.

#### Immunogenicity in AGMs

Evaluation of the immunogenicity of Ad26.RSV.preF and RSV preF protein in RSV pre-exposed AGMs (*Chlorocebus aethiops sabaeus*; Wake Forest School of Medicine, Winston-Salem, NC, USA) was conducted as follows: healthy, nonpregnant, adult and aged female AGMs (9-26 years of age) were housed in 7 different social groups with additional non-study animals. Ultrasounds were performed to monitor pregnancies; body temperature and body weight were measured each time the animals underwent anesthesia. AGMs (n = 42) received i.n. RSV Memphis 37 (7.5×10^5^ pfu; 500 µL in each nostril for a total of 1 mL) and were temporarily removed from their social groups for 2 weeks following RSV inoculation. RSV F–binding antibody titers were measured by ELISA at Week –5; 36 animals were divided into 5 experimental groups, with comparable RSV F ELISA titers, age distributions, and division between social groups. Animals received i.m. immunization with Ad26.RSV.preF alone (1×10^11^ vp [n = 11]), RSV preF protein alone (150 µg [n = 4]), or a combination of Ad26.RSV.preF (1×10^11^ vp) and RSV preF protein (15 µg, 50 µg, or 150 µg; [n = 7 each group]) at Week 0. Animals who showed signs of illness or were pregnant at the time of immunization were removed from the study, and the number of animals throughout the study were as follows: Ad26.RSV.preF (Week 0 to Week 59, n = 11; Week 60 to Week 104, n = 9); RSV preF protein (Week 0 to Week 95, n = 4; Week 96 to Week 104, n = 3); Ad26.RSV.preF/RSV preF protein 150 µg (Week 0 to Week 50, n = 7; Week 51 to Week 59, n = 6; Week 60 to Week 104, n = 4); Ad26.RSV.preF/RSV preF protein 50 µg (Week 0 to Week 59, n = 7; Week 60 to Week 104, n = 6); and Ad26.RSV.preF/RSV preF protein 15 µg (Week 0 to Week 59, n = 7; Week 60 to Week 104, n = 6).

Animals received homologous boost immunization at Week 60. Blood samples were collected throughout the study for serum and peripheral blood mononuclear cell (PBMC) isolation. BAL was collected at Weeks –1 and 2, and nasal swabs were obtained at Weeks –1, 2, 60, 61, and 104.

#### Virus neutralization assays

Virus neutralization assays were performed similarly for naïve and RSV pre-exposed mice, cotton rats, and AGMs. Heat-inactivated animal serum samples were serially diluted and mixed with 25,000 pfu of firefly luciferase (FFL)–labeled RSV CL57 or RSV A2 propagated on A549 cells (American Type Culture Collection, Manassas, VA, USA) in half-area white tissue culture plates and incubated for 1 hour at room temperature (RT). Subsequently, 5×10^3^ A549 cells (multiplicity of infection: 5) per well were added, and plates were incubated for 20 hours at 37°C in 10% CO_2_. After 20 hours, neolite substrate (PerkinElmer, Waltham, MA, USA) was added, and the luminescence signal was measured using an Envision® plate reader (PerkinElmer) or a BioTek Synergy NEO plate reader (Agilent Technologies, Santa Clara, CA, USA); VNT were reported as the 90% inhibitory concentration (IC_90_) for mice and as the 50% inhibitory concentration (IC_50_) for cotton rats and AGMs.

#### ELISA

ELISA was used to measure RSV preF–specific IgG antibodies or IgG1 and IgG2a subclass antibodies in mice. Briefly, 96-well plates were coated with anti–RSV F (1 µg/mL; MedImmune, Gaithersburg, MD, USA) or with streptavidin (0.66 µg/mL) and incubated at 4°C overnight. Wells were washed and then blocked with bovine serum albumin for 30 minutes at RT. After washing, RSV preF protein (0.25 µg/mL, anti–RSV F–coated wells) or biotinylated RSV preF protein (0.3125 µg/mL, streptavidin-coated wells) was added and incubated for 1 hour at RT. Wells were washed again and serially diluted; heat-inactivated serum samples and standards were added to the wells and incubated for 1 to 2 hours at RT. RSV preF–specific antibodies were detected by horseradish peroxidase (HRP)–labeled anti-mouse IgG, IgG1, or IgG2a (Southern Biotech, Birmingham, AL, USA). For anti–RSV F/RSV preF–coated wells, the reaction was developed with O-phenylenediamine dihydrochloride substrate. The reaction was stopped by addition of 1M H_2_SO_4_, and the optical density was measured at 492 nm. The titers were expressed as log_10_ half-maximum effective concentration (EC_50_) titers. For the biotinylated RSV preF protein–coated wells, the reaction was developed with LumiGLO® substrate (SeraCare, Milford, MA, USA) and the luminescence signal was measured at 428 nm. Titers were expressed as log_10_ relative potency (in comparison with a standard serum sample).

In AGMs, RSV preF–specific IgG (in serum and BAL) and RSV preF–specific IgA (in nasal swabs and BAL) titers were determined by ELISA. White 96-well plates were coated with streptavidin (1 µg/mL) and incubated overnight at 4°C; after washing, wells were blocked with fetal bovine serum albumin for 30 minutes at RT. Wells were washed again, followed by the addition of biotinylated RSV preF protein and incubation for 1 hour at RT. After washing, serially diluted heat-inactivated serum samples were added and incubated for 2 hours at RT. HRP-labeled anti-human IgG (Jackson ImmunoResearch Laboratories, Inc., West Grove, PA, USA) or HRP-labeled anti-monkey IgA (Sigma-Aldrich, St. Louis, MO, USA) was added to the wells for detection; the reaction was developed with LumiGLO® substrate and the luminescence signal was measured at 428 nm. For BAL and nasal swabs, total protein was measured by OD280 measurement, which was used to correct for sample concentration. RSV preF–specific IgG-binding titers in serum were expressed as log_10_ EC_50_ titers, and RSV preF–specific IgA- and IgG-binding titers in BAL and nasal swabs were expressed as log_10_ endpoint titers divided by protein concentration (mg/mL).

#### Plaque titration assay for RSV titers in cotton rat challenge model

Lung and nasal tissues were obtained postsacrifice on Day 54. Left lung lobes from each cotton rat were weighed prior to homogenization; nasal tissue sample mass was estimated to be 0.3 g. Homogenized tissue samples (undiluted, 1:10 dilution, and 1:100 dilution) were used for plaque titration assays; each sample was assayed in duplicate. Samples (100 µL) were added to 24-well plates confluent with Hep-2 cells and incubated for 2 hours; supernatant was aspirated, and wells were overlayed with methylcellulose media for 6 days until plaque formation. Plates were stained with crystal violet, and plaques were counted to determine the virus concentration (pfu). Challenge virus concentration was similarly measured immediately after challenge to confirm challenge dose.

#### IFN-γ ELISpot

Mice were sacrificed at Week 5 (RSV pre-exposed mice) or Week 6 (naïve mice); spleens were collected and processed into a single-cell suspension using a gentleMACS™ Dissociator (Miltenyi Biotec, North Rhine-Westphalia, Germany). RSV F–specific IFN-γ ELISpot assays were performed by seeding 500,000 splenocytes per well and stimulating overnight with a peptide pool representing the complete RSV A2 F protein sequence (JPT Peptide Technologies, Berlin, Germany). Spot-forming units (SFU) were quantified using a mouse IFN-γ ELISpot kit (Mabtech, Stockholm, Sweden) per the manufacturer’s instructions. Background levels were determined using the 95^th^ percentile of the SFU in nonstimulated wells.

AGM blood samples were shipped overnight in heparin tubes at RT to Charles River Laboratories. PBMCs were isolated, and cellular responses were determined on fresh PBMCs at Charles River Laboratories using a monkey IFN-γ ELISpot kit (Mabtech) at Weeks 1, 2, 7, 9, 15, and 72; PBMCs (2×10^5^) were tested in triplicate for response to the RSV A2 F peptide pool stimulation (2 µg per peptide/mL). PBMCs (2×10^4^) were tested in duplicate either unstimulated for background measurements or in response to positive control (phytohemagglutinin [PHA]); averaged background (unstimulated) values were subtracted from the averaged stimulated values.

#### ICS

Antigen-specific cellular immune responses were measured by ICS on mouse splenocytes isolated at Week 8. Splenocytes were stimulated overnight with an RSV F peptide pool, hamster–anti-mouse CD28, and rat–anti-mouse CD49d. BD GolgiPlug™ was added after 1 hour. Samples were stained by amine-reactive violet dye (Invitrogen, Waltham, MA, USA) for dead cell discrimination. Anti-mouse CD16/CD32 antibodies were used to block Fc receptors, and cells were stained with anti–CD3-FITC (Clone 142-2C11, BD Biosciences), anti–CD4-PerCpCy5.5 (Clone RM4-5, BD Biosciences), and anti–CD8-APC-H7 (Clone 53-6.7, BD Biosciences). The cells were then permeabilized with BD Cytofix/Cytoperm™ and subsequently intracellularly stained using anti–IFN-γ-PE (Clone XMG1.2), anti–TNF-α-PE-Cy7 (Clone MP6-XT22), and anti–IL-2-APC (Clone JES6-5H4) antibodies. The percentage of CD3^+^CD4^+^ and CD3^+^CD8^+^ T cells expressing IFN-γ, TNF-α, or IL-2 was quantified by flow cytometry using a BD FACSCanto™ II. All reagents were from BD Biosciences, San Diego, CA, USA. Analysis of flow cytometric data was performed in FlowJo software version 9.6.1 (FlowJo, LLC, Ashland, OR, USA).

ICS was performed on AGM PBMCs at Charles River Laboratories. Briefly, 1×10^6^ PBMCs were isolated (as previously described) and tested in triplicate for response to the RSV A2 F peptide pool (2 µg per peptide/mL) or PMA/Ionomycin. Cells were incubated for 1 hour at 37°C, followed by the addition of a BD GolgiStop/GolgiPlug mix solution (BD Biosciences) and further incubation for 5 hours and overnight storage at 4°C. Cells were then stained with a live-dead viability dye (Invitrogen) and the following antibody panel: CD4-FITC (Clone L200, BD Biosciences), IL-2-PE (Clone M7/48A, Miltenyi Biotec), CD14-PerCP (Clone TUK4, Miltenyi Biotec), CD8-PE-Cy7 (Clone SK1, BioLegend, San Diego, CA, USA), IFN-γ-APC (Clone B27, BioLegend), CD3-APC-Cy7 (Clone SP34, BD Biosciences), and TNF-α-BV421 (Clone Mab11, BioLegend); the cells were analyzed using a BD FACSCanto™ II flow cytometer.

#### ADCC reporter assay

A bioluminescent murine, FcγRIV-specific, ADCC reporter assay was used. A549 cells in 96-well culture plates were infected with RSV Long overnight. The cells were washed and serially diluted serum pools (from mice immunized with Ad26.RSV.preF or RSV preF protein) were added, together with Jurkat cells (90,000 cells per well) stably transfected with mouse FcγRIV and NFAT-REluc2 expression plasmids (Promega, Madison, WI, USA). After incubation at 37°C for 6 hours, Bio-Glo™ Luciferase Assay reagent (Promega) was added and luminescence was measured. To control for antibody binding, RSV-infected cells incubated with the serially diluted serum pools were fixed and stained with AF488-labeled goat anti-mouse IgG (Invitrogen).

### Statistical analysis

#### Mouse studies

For studies in mice, analysis of variance (ANOVA) was used for statistical comparisons between groups. For VNT, ELISA, ELISpot, and ICS, difference testing was applied using ANOVA for potentially censored observations; the data were log_2_ transformed (VNT, ELISA) or log_10_ transformed (ELISpot and ICS). In the RSV-naïve mouse study, a stepwise approach for difference testing was applied, starting with the highest RSV preF protein dose and analyzing the next lower dose if a significant difference was observed. In the RSV pre-exposed mouse study, an across–Ad26.RSV.preF dose analysis was done. Statistical analyses were performed using SAS version 9.4 (SAS Institute, Inc., Cary, NC, USA).

#### Cotton rat studies

Lung and nasal tissue viral titers in cotton rats were compared using a Cochran-Mantel-Haenszel test based on rank scores with the Ad26.RSV.preF dose as the stratification variable. Comparisons were made between groups receiving Ad26.RSV.preF alone versus those receiving Ad26.RSV.preF/RSV preF protein combinations. Exploratory comparisons between animals receiving Ad26.RSV.preF/RSV preF protein combinations versus those receiving RSV preF protein alone were made using a Wilcoxon rank sum test with Bonferroni correction.

#### AGM studies

For studies in AGMs, comparisons between groups receiving Ad26.RSV.preF alone, RSV preF protein alone, or Ad26.RSV.preF/RSV preF protein combinations were made per readout per time point using ANOVA for potentially censored measurements (Tobit model) and included a Dunnett correction for multiple testing. Similarly, changes over time within groups (before and after prime immunization; or before and after boost immunization) were analyzed based on a paired *t* test with a Bonferroni correction.

## Data availability

All data to understand and assess the conclusions of this research are available in the main text and **Supplementary Materials**. The raw data that support the findings of this study are available from the corresponding author upon reasonable request.

## Acknowledgments

This study was funded by Janssen Pharmaceuticals. Medical writing support was provided by William J. Kelley, PhD (Lumanity Communications Inc., Yardley, PA, USA), and was funded by Janssen Global Medical Affairs. The AGM colony at Wake Forest was supported by NIH grant P40-OD010965.

## Author contributions

E.S., L.v.d.F, F.C., and R.C.Z. designed the experiments, analyzed the data, and wrote the manuscript. J.P.M.L, H.S., B.C., and R.C.Z. contributed to the conception of the work. R.B., M.H.-v.d.M., J.D., Y.v.P., C.V., and L.T. performed the experiments and analyzed the data. J.S. performed the statistical analyses. M.J.J. supervised all *in vivo* AGM procedures. All authors reviewed, critiqued, and approved the text.

