## Supplementary Figures 1, 2, and 3 for "Combination Ad26.RSV.preF/RSV preF protein vaccine induces superior humoral, cellular, and protective immunity compared with individual vaccine components in preclinical models"

**Supplementary Materials**

**Supplementary Figure 1. ICS measured at Week 5 in splenocytes isolated from RSV pre-exposed mice.** Female BALB/c mice were pre-exposed i.n. with RSV A2 ($5\times10^5$ pfu) at Week –79 and immunized intramuscularly at Week 0 with Ad26.RSV.preF ($1\times10^8$ or $1\times10^9$ vp; n = 12 in each group), RSV preF protein (0.15 µg; n = 6), Ad26.RSV.preF ($1\times10^8$ or $1\times10^9$ vp) combined with RSV preF protein (0.15 µg; n = 12 each group). Negative control animals received formulation buffer (n = 6). Cellular responses were measured by ICS in splenocytes at Week 5, and the percentage of CD4+ (left panels) and CD8+ (right panels) T cells expressing IFN-γ (top panels), TNF-α (middle panels), and IL-2 (bottom panels) is shown. The horizontal lines indicate the geometric mean titer per group, and the dotted line indicates 3×SD above the mean response of unstimulated cells.

Ad26, adenovirus type 26; ICS, intracellular cytokine staining; IFN-γ, interferon-γ; IL-2, interleukin-2; i.n., intranasally; ns, not significant; pfu, plaque-forming units; preF, prefusion conformation–stabilized RSV F protein; RSV, respiratory syncytial virus; SD, standard deviation TNF-α, tumor necrosis factor α; vp, viral particles.

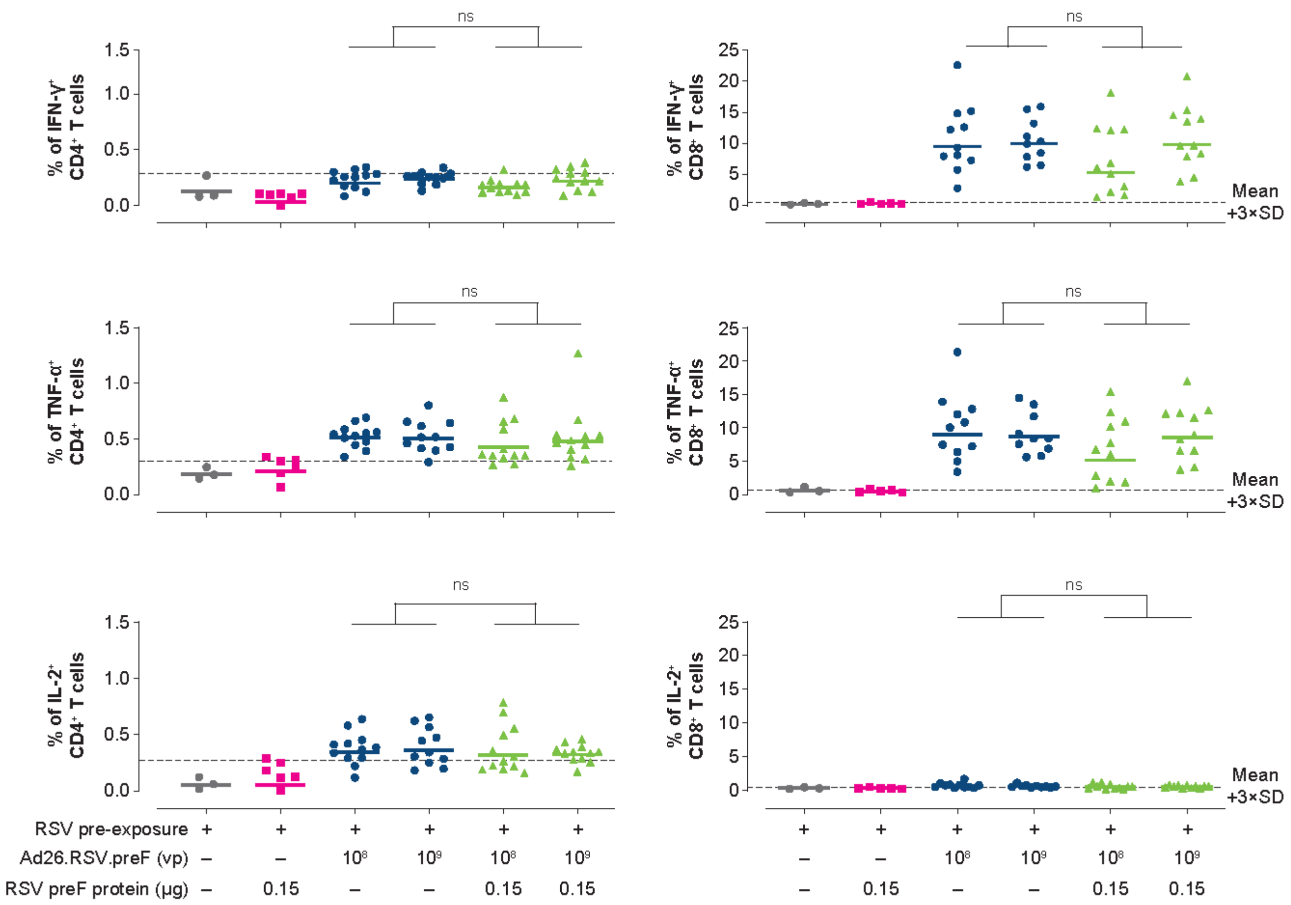
ns
% of IFN-γ+ CD4+ T cells
% of IFN-γ+ CD8+ T cells
Mean +3×SD
% of TNF-α+ CD4+ T cells
% of TNF-α+ CD8+ T cells
% of IL-2+ CD4+ T cells
% of IL-2+ CD8+ T cells
RSV pre-exposure
Ad26.RSV.preF (vp)
RSV preF protein (µg)
0.15

**Supplementary Figure 2. ICS measured from PBMCs isolated from African green monkeys.** Female African green monkeys were pre-exposed i.n. with RSV Memphis 37 ($7.5\times10^5$ pfu) at Week –19 and immunized intramuscularly with Ad26.RSV.preF ($1\times10^{11}$ vp; n = 11); RSV preF protein (150 µg; n = 4); or a combination of Ad26.RSV.preF ($1\times10^{11}$ vp) with RSV preF protein (150 µg; n = 7), RSV preF protein (50 µg; n = 7), or RSV preF protein (15 µg; n = 7) at Week 0. PBMCs were obtained before immunization and after immunization at Weeks 2, 7, and 9 for measurements of cellular immune responses by ICS. The percentage of CD8+ T cells expressing IFN-γ, TNF-α, and IL-2 in PBMCs of individual animals is shown. Horizontal bars indicate geometric mean responses.

Ad26, adenovirus type 26; ICS, intracellular cytokine staining; IFN-γ, interferon-γ; IL-2, interleukin-2; i.n., intranasally; PBMC, peripheral blood mononuclear cell; pfu, plaque-forming units; preF, prefusion conformation–stabilized RSV F protein; RSV, respiratory syncytial virus; TNF-α, tumor necrosis factor α; vp, viral particles.

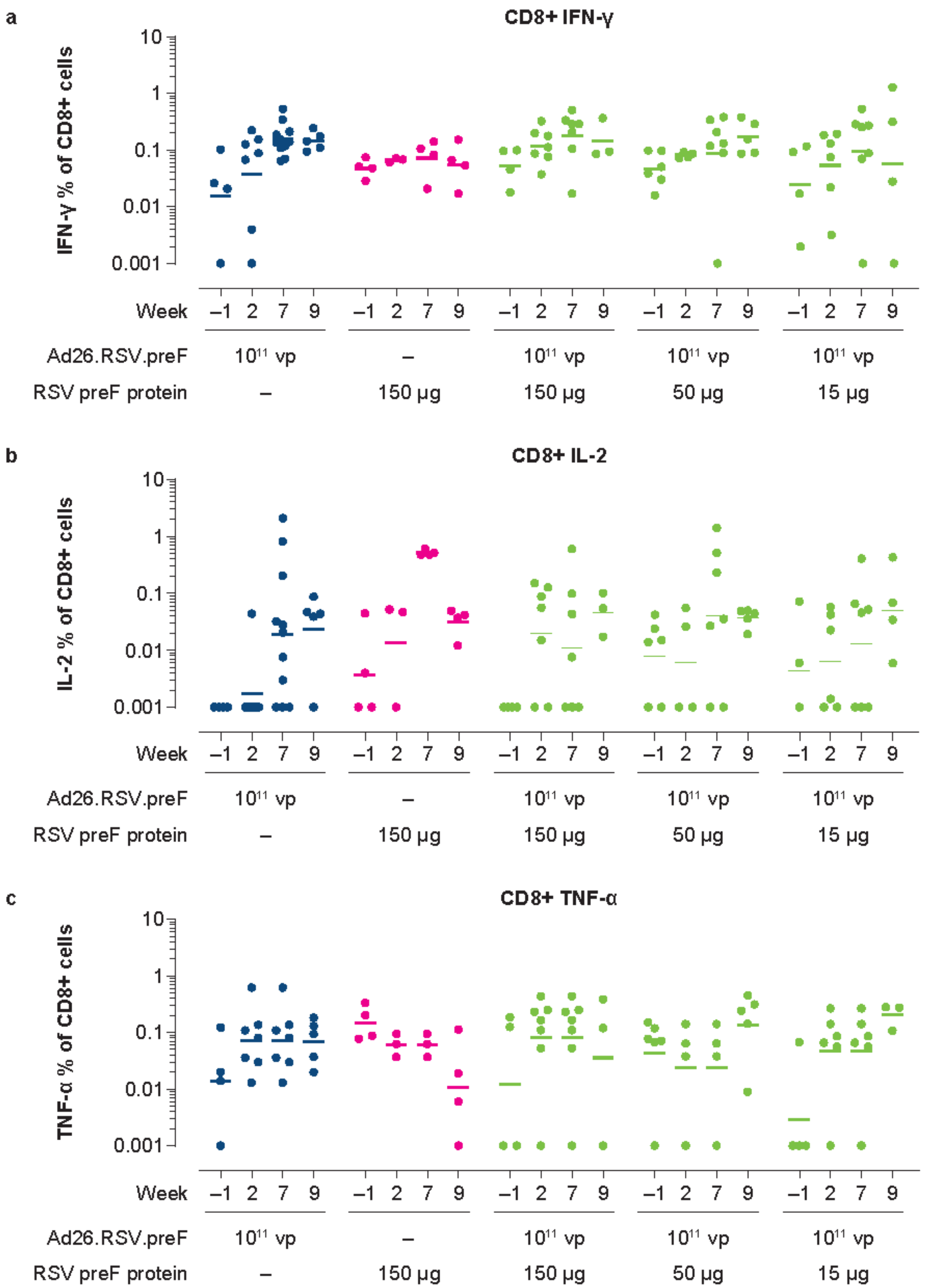

a
CD8+ IFN-γ
IFN-γ % of CD8+ cells
10
1
0.1
0.01
0.001
b
CD8+ IL-2
IL-2 % of CD8+ cells
c
CD8+ TNF-α
TNF-α % of CD8+ cells
Week –1 2 7 9 –1 2 7 9 –1 2 7 9 –1 2 7 9 –1 2 7 9
Ad26.RSV.preF 10^11 vp – 10^11 vp 10^11 vp 10^11 vp
RSV preF protein – 150 µg 150 µg 50 µg 15 µg

**Supplementary Figure 3. ICS measured from PBMCs isolated from African green monkeys.** Female African green monkeys were pre-exposed i.n. with RSV Memphis 37 ($7.5\times10^5$ pfu) at Week –19 and immunized intramuscularly with Ad26.RSV.preF ($1\times10^{11}$ vp; n = 11); RSV preF protein (150 µg; n = 4); or a combination of Ad26.RSV.preF ($1\times10^{11}$ vp) with RSV preF protein (150 µg; n = 7), RSV preF protein (50 µg; n = 7), or RSV preF protein (15 µg; n = 7) at Week 0. PBMCs were obtained before immunization and after immunization at Weeks 2, 7, and 9 for measurements of cellular immune responses by ICS. The percentage of CD4+ T cells expressing IFN-γ, TNF-α, and IL-2 in PBMCs of individual animals is shown. Horizontal bars indicate geometric mean responses.

Ad26, adenovirus type 26; ICS, intracellular cytokine staining; IFN-γ, interferon-γ; IL-2, interleukin-2; i.n., intranasally; PBMC, peripheral blood mononuclear cell; pfu, plaque-forming units; preF, prefusion conformation–stabilized RSV F protein; RSV, respiratory syncytial virus; TNF-α, tumor necrosis factor α; vp, viral particles.

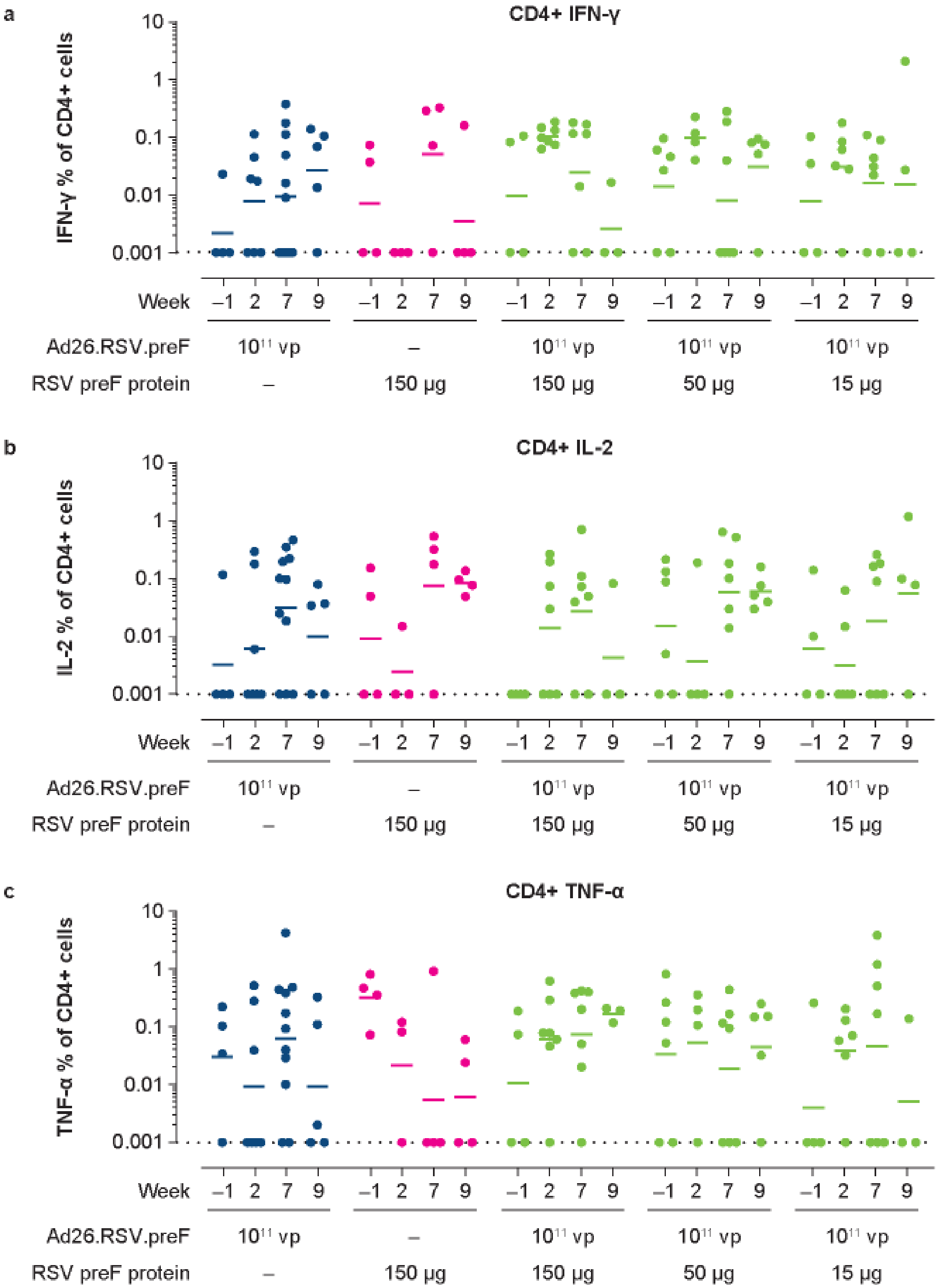
a
CD4+ IFN-γ
IFN-γ % of CD4+ cells
10
1
0.1
0.01
0.001
Week –1 2 7 9 –1 2 7 9 –1 2 7 9 –1 2 7 9 –1 2 7 9
Ad26.RSV.preF 10[11] vp – 10[11] vp 10[11] vp 10[11] vp
RSV preF protein – 150 μg 150 μg 50 μg 15 μg
b
CD4+ IL-2
IL-2 % of CD4+ cells
10
1
0.1
0.01
0.001
Week –1 2 7 9 –1 2 7 9 –1 2 7 9 –1 2 7 9 –1 2 7 9
Ad26.RSV.preF 10[11] vp – 10[11] vp 10[11] vp 10[11] vp
RSV preF protein – 150 μg 150 μg 50 μg 15 μg
c
CD4+ TNF-α
TNF-α % of CD4+ cells
10
1
0.1
0.01
0.001
Week –1 2 7 9 –1 2 7 9 –1 2 7 9 –1 2 7 9 –1 2 7 9
Ad26.RSV.preF 10[11] vp – 10[11] vp 10[11] vp 10[11] vp
RSV preF protein – 150 μg 150 μg 50 μg 15 μg
